# How epigenome drives chromatin folding and dynamics, insights from efficient coarse-grained models of chromosomes

**DOI:** 10.1101/200584

**Authors:** Surya K. Ghosh, Daniel Jost

## Abstract

The 3D organization of chromosome is crucial for regulating gene expression and cell function. Many experimental and polymer modeling efforts are dedicated to deciphering the mechanistic principles behind chromosome folding. Chromosomes are long and densely packed - topologically constrained - polymers. The main challenge is therefore to develop adequate models and simulation methods to investigate properly the multi spatio-temporal scales of such macromolecules. Here, we discussed a generic strategy to develop efficient coarse-grained models for self-avoiding polymers on a lattice. Accounting accurately for the polymer entanglement length and the volumic density, we show that our simulation scheme not only captures the steady-state structural and dynamical properties of the system but also tracks the same dynamics at different coarse-graining. This strategy allows a strong power-law gain in numerical efficiency and offers a systematic way to define reliable coarse-grained null models for chromosomes and to go beyond the current limitations by studying long chromosomes during an extended time period with good statistics. We use our formalism to investigate in details the time evolution of the 3D organization of chromosome 3R (20 Mbp) in drosophila during one cell cycle (20 hours). We show that a combination of our coarse-graining strategy with a one-parameter block copolymer model integrating epigenomic-driven interactions quantitatively reproduce experimental data at the chromosome-scale and predict that chromatin motion is very dynamic during the cell cycle.

## 1 Introduction

Though all cells of a multicellular organism contain the same genetic information, they vary widely in shapes, in physiologies, and in functions. These differences mainly reflect variations in gene expression between different tissues or cell types. Recent experiments have highlighted the important role of the physical organization of chromosomes inside the cell nucleus in regulating gene expression^1–3^: gene activities being modulated, not only by the local folding of the chromatin fiber but also by its higher order organization with 3D nuclear compartments favorable to gene activation or repression. During interphase, the longest phase of the cell cycle where genes are expressed and DNA is replicated, chromosomes are found to be organized hierarchically. Confocal and electron microscopy experiments have revealed that each chromosome occupies it own territory^4^. Also, the genes sharing the same transcriptional state tend to colocalize^5–7^: inactive genomic regions (the heterochromatin) being more peripheral while active regions (the euchromatin) being more central. At the sub-chromosomal level, advanced molecular biology tools, like chromosome conformation capture (Hi-C) techniques, have shown that chromosomes are partitioned into consecutive 3D interaction compartments, the so-called topologically-associated domains (TADs),^8–10^. Loci inside these domains experience enriched contact probabilities with other loci of the same domain while showing partial insulation from loci of nearest neighbor domains. These domains can be easily visualized as consecutive “squares” along the diagonal of a 2D contact frequency matrix (see Fig.6 for an illustration). TADs formation has been associated with the local biochemical composition of chromatin, the so-called epigenome, which encodes for gene activity^7,11–14^: genes inside the same TAD tends to have the same epigenomic state, and long-range contacts may be observed between TADs of the same state.

However, how genome precisely organized in space is still not fully understood and addressing this question represents one of the most exciting challenges of modern biology^15^. Lots of experimental and modeling efforts are currently dedicated to understanding the mechanisms implied in chromosome folding. In particular, polymer models have been instrumental in simulating and testing different molecular and physical mechanisms and in driving new experiments^5,16–39^. An important challenge for such models is to simulate with a good precision the behavior of long polymer chains (the typical size of a chromosome ranging from about a million base-pairs in yeast to hundreds of Mbp in human) during an extended time period (of the order of hours for a typical cell cycle). Therefore the standard strategy used in these approaches is to start from a *coarse-grained “null”* model for chromatin with few basic interactions^40^. Then eventually decorate it with more physical or chemical interactions driven by biological information such as the gene activity or the local epigenomic state.

Chromosomes being very long polymers densely packed into the cell nucleus, topological constraints generated by polymer entanglement may play an important role in controlling the dynamics and fluctuations of such polymers^16,41^. However, when building their null models, very few approaches account adequately for such considerations. As neglecting topological constraints may lead to different structural and dynamical properties of the polymer, how can one interpret the outcomes correctly of such models concerning realistic mechanisms, if the considered null model is already biased?

Here, we develop a generic coarse-graining strategy for self-avoiding polymers that allows, simultaneously, to drastically reduce the computation time while maintaining the polymer in the same topological regime and thus preserving the correct structural and dynamical properties. In the first part, we explain the strategy and demonstrate its efficiency, leading to a systematic approach to developing a coarse-grained null model for chromosomes. In the second part, we apply it to investigate the role of epigenomic-driven interactions in the folding and dynamics of drosophila chromosomes. Finally, we discuss our results and their implications in the general context of chromosome modeling and chromatin biology.

## 2 Results

### 2.1 Chromosome and entanglement length

Chromosomes are long polymers confined inside a small volume, the cell nucleus^42^. As a result, the generic characteristics of these densely packed long polymers are very different from free isolated chains and exhibit distinct universal properties^41,43^. For a simple semi-flexible self-avoiding polymer, composed of *N* beads of size *b* (in *nm*), each bead representing *v bp*, such properties are mainly determined by (i) its rigidity, characterized by its Kuhn length *l_k_* (in *nm*), and (ii) by its volumic density *ρ* (in *bp/nm*^3^). Moreover, in a non-dilute environment, topological constraints are also expected to influence the large-scale organization and long-time dynamics. Their importance depends on the ratio between the polymer contour length *L* ≡ *Nb* and *L_e_* (in *nm*), the so-called entanglement length. *L_e_* measures the typical subchain size above which topological confinement due to excluded volume influences configurational fluctuations, and depends on *l_k_* and *ρ*. It may be associated with the tube diameter in the reptation model or to the crossover time between a Rouse-like motion and a reptation-like motion^44^, and may be estimated using the phenomenological relation^45^

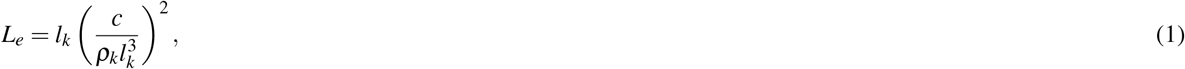

with *c* ≈ 19 is a numerical factor and *ρ_k_* = (*ρ/v*)(*b/l_k_*) the volumic density in Kuhn segment. In the following, we define *N_k_* = *vl_k_/b* (in *bp*) as the Kuhn segment size representing the number of *bp* in one Kuhn segment. *L/L_e_* ≪ 1 means very weak topological effects and the polymer behaves as a standard Rouse chain. If *L/L_e_* ≫ 1, the chain motion is restricted due to strong topological interactions and exploration of the available space is very slow. In this case, equilibration time of the chain scales as *N*^3^^46^, implying that polymer dynamics remains out-of-equilibrium and the initial topological properties (presence/absence of knots) or large-scale organization features are maintained over a long time period.

#### 2.1.1 A reference model for chromosome

To provide the physically realistic scenario of chromosome structure and dynamics, we need to precise the values of *l_k_* and *b*. We define the fine scale null model of chromosomes with Kuhn length 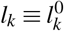, number of beads *N* = No and bead size *b* ≡ *b*_0_ as our reference model. Precise measurements of the Kuhn length of *in vivo* chromatin are still lacking and controversy still exists about its value, going from few nanometers^47^, the so-called 10 *nm*-fiber, to hundreds of nanometers, the so-called 30 *nm*-fiber^48^. We decide to use, as a reference model, the nucleosomal scale (1 monomer correspond to *v*_0_ = 200 bp, *b*_0_ ≈ 10 *nm*) with a recent estimation of Kuhn size 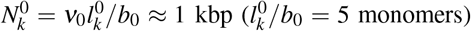 based on cyclization probabilities of chromatin^26^.

In order to determine the value of the entanglement length *L_e_*, the other important quantity to fix is the chromatin volumic density *ρ* defined as the ratio between the genome size and the volume of the nucleus. Depending on the species, the cell types or the developmental stages, it may strongly vary. Typical orders of magnitude are *ρ* ≈ 0.005 *bp/nm*^3^ for haploid yeast, *ρ* ≈ 0.009 *bp/nm*^3^ for drosophila late embryos or *ρ* ≈ 0.015 *bp/nm*^3^ in typical mammalian nuclei (see Materials and Methods). Systems with higher volumic density become more entangled and exhibit shorter entanglement length (Eq. 1). This leads to decreasing values for *N_e_* ≡ *v*_0_*L_e_/b*_0_ ≈ 920 kbp (yeast), 285 kbp (drosophila) or 102 kbp (mammals). In yeast, *v*_0_*N*_0_ ≈ 750 kbp, implying that chromosomes are weakly entangled (*L/L_e_* = 0.8 ≲ 1). In higher eukaryotes, as drosophila or mammals, chromosomes are longer (tens or hundreds of Mbp) and in the regime of strong topological constraints (*L/L_e_* ≫ 1).

For example, for a chromosome of length *v*_0_*N*_0_ = 20 Mbp, the corresponding value is *L/L_e_* = 70 in drosophila. For a given species, the exact value of this ratio may vary depending on the cell types due to variation in the volume of the nucleus but usually the entanglement regime is preserved (weakly constrained for yeast, strongly for higher eukaryotes). Note that the 30 *nm* fiber model for chromatin 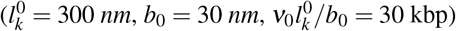 would lead to similar orders of magnitude for *L/L_e_*.

#### 2.1.2 Generic behavior of chromosome

To illustrate the generic behavior of the reference model in the different entanglement regimes, we perform kinetic Monte-Carlo (KMC) simulations of the chain dynamics using a lattice model with periodic boundary conditions and starting from knotted-free initial configurations (see Materials and Methods). We focus on the “yeast” (*N*_0_ = 3750 monomers, 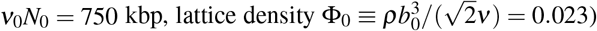) and on the “drosophila” (*N*_0_ = 10^5^ monomers, *ν*_0_*N*_0_ = 20 Mbp, Φ_0_ = 0.043) cases.

During the simulations, we measured four standard physical quantities: time evolution of the average mean squared displacements (MSD) of individual monomers *g*_1_ (*t*), MSD of the center of mass of the chain *g*_3_(*t*), average mean squared distance 〈*R*^2^(*s*)〉 between two monomers separated by a contour length *s* (in *bp*) along the chain and contact probability *P_c_*(*s*) (Fig.1). Comparison of *g*_1_(*t*) with 0.01 *t*^0.5^, the experimentally measured typical value of *g*_1_ in *μm*^2^ for yeast and drosophila^47,49^, gave the equivalency of each *MCS* with real time in *sec*. From the time mapping we were able to represent our results in real physical unit: time in sec, distance in *μm*. For the drosophila case, a 10^8^-MCS long trajectory would correspond to about 30 *min* of real time. To check the precision of the MSD scaling laws, we calculated *g*_1_, *g*_3_ by varying the measuring simulation time window (Δ*t*) of the trajectory. We observed that both *g*_1_, *g*_3_ reached steady-state rapidly and almost perfectly overlap for different trace-lengths Δ*t*, see Fig.S2.

**Figure 1.**
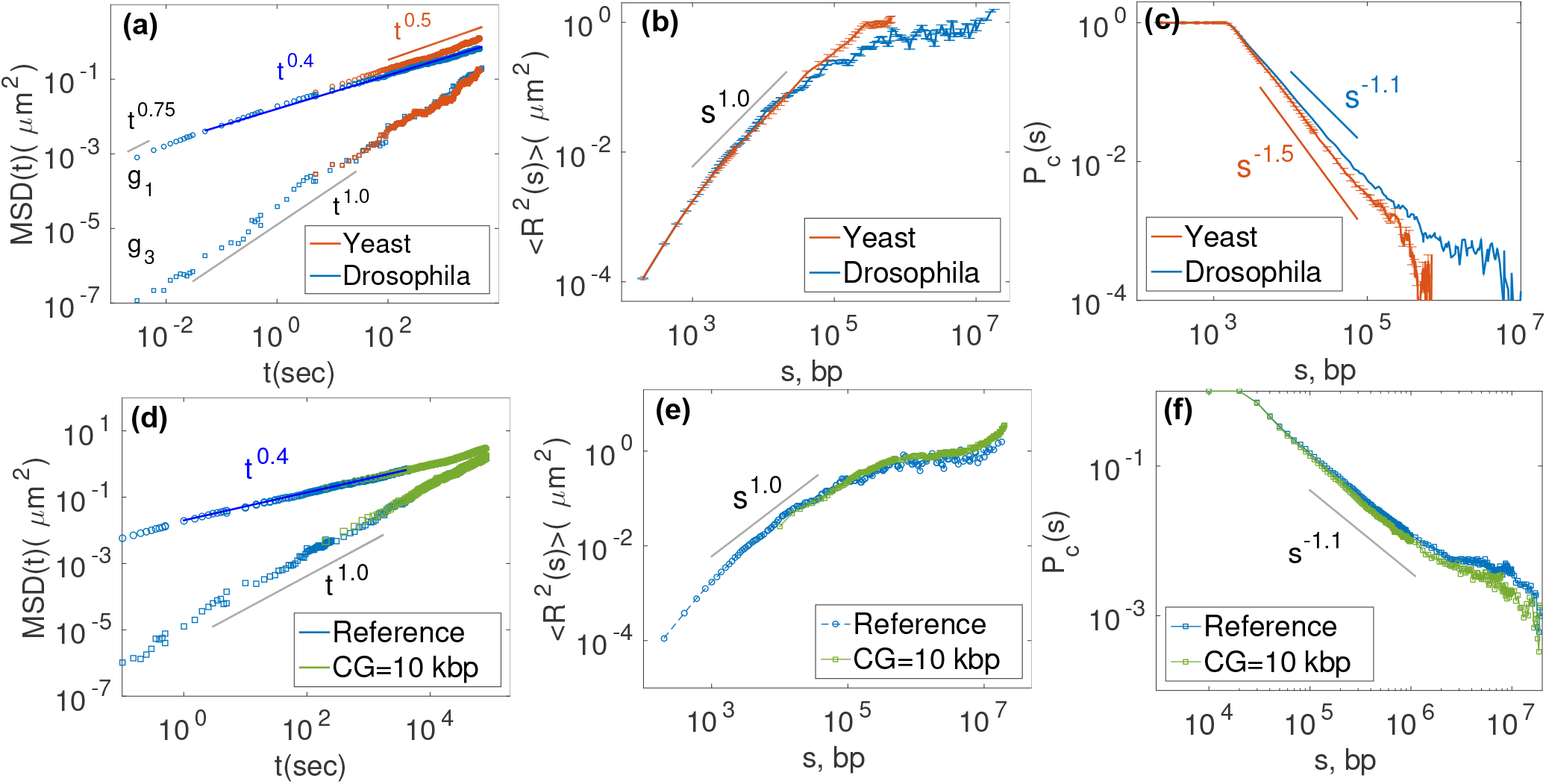
In the top panels, we compare different physical properties for yeast (red) and drosophila (blue) chromosomes at a nucleosomal resolution of 200 *bp* (reference model). In the bottom panel, we compare the reference model (Φ_0_ = 0.043) with one possible coarse-grained model (*CG* = 10 kbp, Φ = 0.97) for the drosophila case. (a,d) Individual MSD *g*_1_ (*t*) (top curves), and center of mass MSD *g*_3_ *(t)* (bottom curves) as a function of time *t*. (b,e) The average physical squared distance *〈R^2^(s)〉* between any two monomers as a function of their linear distance *s* along the chain (given in *bp*). (c,f) Average contact probability *P_c_(s)* as a function of *s*. A contact between any two monomers is defined if the 3D distance is less than a threshold *d_c_* (with *d_c_* = 55 *nm* in (c) and *d_c_* = 163 *nm* in (f)). In (e,f), averages were computed over the same real time window (100 *sec* - 30 *min*). The error bars in (a, b, c) were computed as the standard deviation of the mean. Error bars in (a) are of the similar size of the symbols. For the yeast case, we fix *L/L_e_* = 0.8, *p* = 0.005*bp/nm*^3^, and for drosophila, *L/L_e_* = 70, *p* = 0.009*bp/nm*^3^ (see section 2.1.1).

The yeast chromosome behaves dynamically as a standard Rouse chain^47^. At intermediate times, *g*_1_ ~ *t*^0.5^ (Fig.1a), the typical scaling law in the Rouse diffusion regime^50^. At later time, when *t* is greater than the Rouse time, the typical time by which the polymer has already traveled a distance equivalent to its size, *g*_1_ coincides with the center of mass MSD (*g*_3_ ~ *t*), characteristic of a simple diffusion process^50^. In the drosophila case, topological constraints are strong and the anomalous diffusion exponents of *g*_1_ at intermediate time scale behaves as *t*^0.4^ (Fig.1a,d). Note that, we do not observe the scaling exponent, at least in the scanned time scales, expected from reptation dynamics (*t*^0.25^) of entangled polymers^46^. This is a characteristic of knot-free polymers, like crumpled or ring polymers^51^ and is reminiscent of the initial unknotted configurations. Starting from random configurations that contain complex knots (Fig.S1g), we recover the standard reptation regime (Fig.S4a,d).

At small time scales (*t* < *ms*), *g*_1_ scales as *t*^0.75^ which corresponds to the typical diffusion regime of a semi-flexible chain up to the Kuhn length scale^52^.

Regarding the structural properties 〈*R*^2^(*s*)〉 (Fig.1b) and *P_c_*(*s*) (Fig.1c), we recover the main scaling laws observed experimentally for chromosomes of yeast, fly and other eukaryotes^5,6,43,53–55^. The yeast case is fully consistent with a worm-like-chain at equilibrium with 〈*R*^2^(*s*)〉 ~ *s*^1^ and *P_c_* ~ *s*^−1.5^^50^. On the other hand, for the fly chromosome, the scaling laws (〈*R*^2^(*s*)〉 ~ *s*^2/3^ and *P_c_* ~ *s*^−11^) are consistent with crumpled polymer physics^16,43,56–58^. The large scale behavior (*s* > 1 Mbp) is a remaining signature of the initial scaling laws (Fig.S7): the system has yet to reach steady-state and is still strongly out-of-equilibrium.

### 2.2 Coarse-graining long polymers at fixed entanglement length

Using the fine scale reference model, we recover the expected structural and dynamical behaviors in the different entanglement regimes, fully consistent with previous theoretical works on knot-free and crumpled polymers^16,43,51,56,57,59^ and with experiments^5,6,47,49,53–55,60^. At this nucleosomal scale, the model has very good spatial (10 *nm)* and temporal (15 *μsec*, 30 *min* ≈ 10^8^ MCS) resolutions. However, the underlying cost of this is a prohibitive computational time. For example, for long chromosomes such as in the drosophila case, simulating one 30 *min* long trajectory requires 83 hrs CPU time on a 3.20 GHz CPU. To access more biologically-relevant time-scales (dozens of hours) with good statistics, we aim to develop a coarse-graining strategy of the reference model that allows to speed up the simulation of long trajectories while preserving the main physical characteristics of the original -fine scale - model.

#### 2.2.1 Coarse-graining strategy

We consider an arbitrary fine-scale model (FSM) of a semi-flexible self-avoiding walk defined by 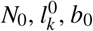. We note *N, l_k_* and *b*, the corresponding values of a coarse-grained model (CGM) of the FSM. Each CGM monomer encompasses *n* = *N*_0_/*N* > 1 FSM monomers. A possible CG strategy consists in neglecting the bending rigidity (*l_k_* = *b*) in the CGM if *n* is greater than the Kuhn size 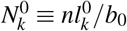, in the FSM, and in imposing the size of CGM monomer to equal the mean end-to-end distance of the corresponding number of FSM monomers, ie 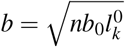. Using Eq.1 and conservation of total volume, it is easy to show that the ratio (*L/L_e_*) is also conserved, and therefore the effect of topological constraints. However, such approach is limited by the volume fraction Φ occupied by the CG chain (for a lattice model Φ ≡ *N/N_s_* with *N_s_* the number of lattice sites).

Indeed, for lattice or off-lattice (assuming spherical shape for monomer in the FSM and CGM) models

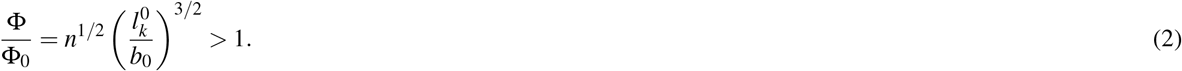

Hence, if Φ_0_ is already high in the FSM and/or the coarse-graining is strong (*n* ≫ 1), Φ might become close or higher to 1 and therefore very hard to simulate. For example in the case of drosophila chromosomes (Φ_0_ ~ 4.3%), if *n* = 5 (corresponding to 1 kbp resolution, the Kuhn segment size of the FSM), Φ = 25Φ_0_ > 1. Therefore, already at the Kuhn size scale, such CG strategy may fail to generate simulable models. Forcing the CG to high *n* anyway would imply to choose 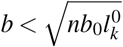 in order to maintain Φ < 1, violating the conservation of the ratio *L/L_e_.* This may affect the dynamical regime of the chain, and hence its physical properties (see Section 2.3). To avoid this, we develop a novel coarse-graining strategy that allows to go for high coarse-graining while keeping the volumic fraction in a simulable range and the ratio *(L/L_e_)* fixed.

We authorized the CGM, even if 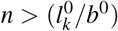, to have a bending rigidity characterized by *l_k_*. And we imposed that the ratio *L/L_e_* and the volume of the simulation box are conserved. In the lattice framework, using Eq. 1, these constraints can be reformulated as (see Materials and Methods)

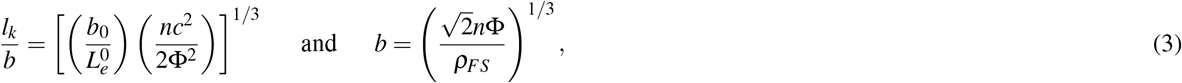

with *ρ_FS_* ≡ *N*_0_/*V* = *ρ/v*_0_ the volumic density in FS monomers (with *V* the volume of the box). Φ now plays the role of a control parameter: the characteristics of the CGM depends not only on the FS properties but also on Φ (see Table 1 for examples) since a given Φ determines *b* and *l_k_*, and the corresponding value for the lattice bending energy *κ* is inferred from *l_k_/b* (see Materials and Methods). As in most coarse-graining approaches, the size of each CG monomer (*b*) does not reflect the actual contour length of the corresponding fine-scale subchain, but rather would represent the typical diameter of the volume occupied by the fine-scale monomers. However, we observe that the length deformation of the CG polymer with respect to the reference model remains weak (Fig.S16a).

**Table 1.** Simulation parameters at different coarse-graining for a semi-flexible, self-avoiding polymer with *L/L_e_* = 70 and *ρ* = 0.009 *bp/nm*^3^ (drosophila case). Lattice volumic fraction Φ, Kuhn size *N_k_* = *vl_k_/b* (in kbp), bond length of the polymer *b* (in *nm*) and Kuhn length *l_k_* (in *nm*) at different coarse-grainings (CG) of *v* = 0.2, 2, 5, and 10 kbp. Each fifth subcolumn represents the time in *msec* equivalent to one Monte-Carlo steps (MCS). Similar tables for the yeast and mammalian cases are given in the supplementary text.

| 200 bp |  |  |  |  | 2 kbp |  |  |  |  | 5 kbp |  |  |  |  | 10 kbp |  |  |  |  |
| --- | --- | --- | --- | --- | --- | --- | --- | --- | --- | --- | --- | --- | --- | --- | --- | --- | --- | --- | --- |
| $\Phi_0$ | $N_k^0$ | $b_0$ | $l_k^0$ | MCS | $\Phi$ | $N_k$ | $b$ | $l_k$ | MCS | $\Phi$ | $N_k$ | $b$ | $l_k$ | MCS | $\Phi$ | $N_k$ | $b$ | $l_k$ | MCS |
| 0.043 | 1.0 | 10.6 | 55.4 | 0.015 | 0.027 | 29 | 20.5 | 305 | 0.002 | 0.09 | 44 | 41.9 | 374 | 0.4 | 0.37 | 44 | 83.8 | 372 | 8.71 |
|  |  |  |  |  | 0.039 | 23 | 23.1 | 273 | 0.010 | 0.17 | 29 | 51.3 | 304 | 1.2 | 0.68 | 29 | 102.5 | 304 | 17.3 |
|  |  |  |  |  | 0.049 | 20 | 24.9 | 252 | 0.040 | 0.24 | 23 | 57.6 | 271 | 1.7 | 0.97 | 23 | 115.3 | 274 | 22.1 |
|  |  |  |  |  | 0.076 | 15 | 28.8 | 218 | 0.083 | 0.29 | 20 | 61.5 | 252 | 2.5 |  |  |  |  |  |
|  |  |  |  |  | 0.113 | 11 | 32.9 | 190 | 0.181 | 0.45 | 15 | 70.9 | 220 | 5.4 |  |  |  |  |  |

It has to be noted that the corresponding CG bending rigidity is not a “true” rigidity that reflects the rigidity of the FSM. It is an artificial rigidity, allowing to control Φ. Therefore, the CGM cannot pretend to quantitatively describe the FSM properties at scales smaller than few *l_k_*.

#### 2.2.2 Conservation of generic properties and time mapping

In this part, we test if the above coarse-graining strategy conserves the structural and dynamical properties of the reference fine-scale model. In this article, we primarily focus on the drosophila case. However, in the Supplementary Information, we show that the method also performs very well for the yeast and mammalian cases (Fig.S8, S9) and that the success of the strategy does not depend on the type of used initial conditions (Fig.S3,S5).

In Fig.1 bottom panel, we compared results between the FSM and a CGM at 10 kbp resolution for Φ = 0.97. Like for the reference model (see Sec.2.1.2), we time-mapped the simulation time for the CGM using *g*_1_(*t*) in order to have a direct time correspondence between the FSM and CGM (see Table 1). For this CG, 1 MCS represents a time step 10^4^ fold larger than the FSM, meaning that the 10^8^ MCS-long trajectories can span more than 100 hours (instead of 30 *min* for the FSM). We remark that MSD curves overlap nicely and that the scaling laws are conserved (Fig.1d). From the configurations collected during the “real” time window (0 - 30 *min*) common to both models, we calculated the averaged properties of 〈*R*^2^〉 (*s*) and *P_c_(s)* (Fig.1e,f). For *s* < 1 Mbp, we observe a perfect match between CG predictions and the FSM behavior. For *s* > 1M*b*, the system does not reach steady-state and keeps a partial memory of the initial scaling laws that are different for both systems (Fig.S6, S7), leading to small deviations between the predictions, especially for *P_c_* which is more sensitive to local structures.

In Fig.2, we performed similar comparisons between two different CGM at a fixed Kuhn size value *N_K_* ≡ *νl_k_/b* (*N_K_* = 23 kbp): *(CG* = 5 kbp, Φ = 0.24) and *(CG* = 10 kbp, Φ = 0.97). As before, we recovered identical scaling laws for *g*_1_(*t*), the 10 kbp-resolution model allowing to scan longer times for the same number of MCS steps (Fig.2a). We computed 〈*R*^2^〉 and *P_c_* for a series of snapshots taken at several real time points, at 1 *min*, 30 *min* and 10 *hrs* (Fig.2b,c). Remarkably, starting from similar behaviors for 〈*R*^2^(*s*)〉 for the two CG (compare the 1 *min*-curves), the predicted time-evolution of the two curves remains identical, even after simulating more than 10h of real time. A similar comparison is also observed for *P_c_(s)* and the average second moment *〈σ^2^(s)〉* of the squared distance *R^2^(s)* defined as *σ^2^(s)* = 〈*R*^4^(s)〉 – 〈*R*^2^(*s*)〉^2^ (Fig.S10c,d).

**Figure 2.**
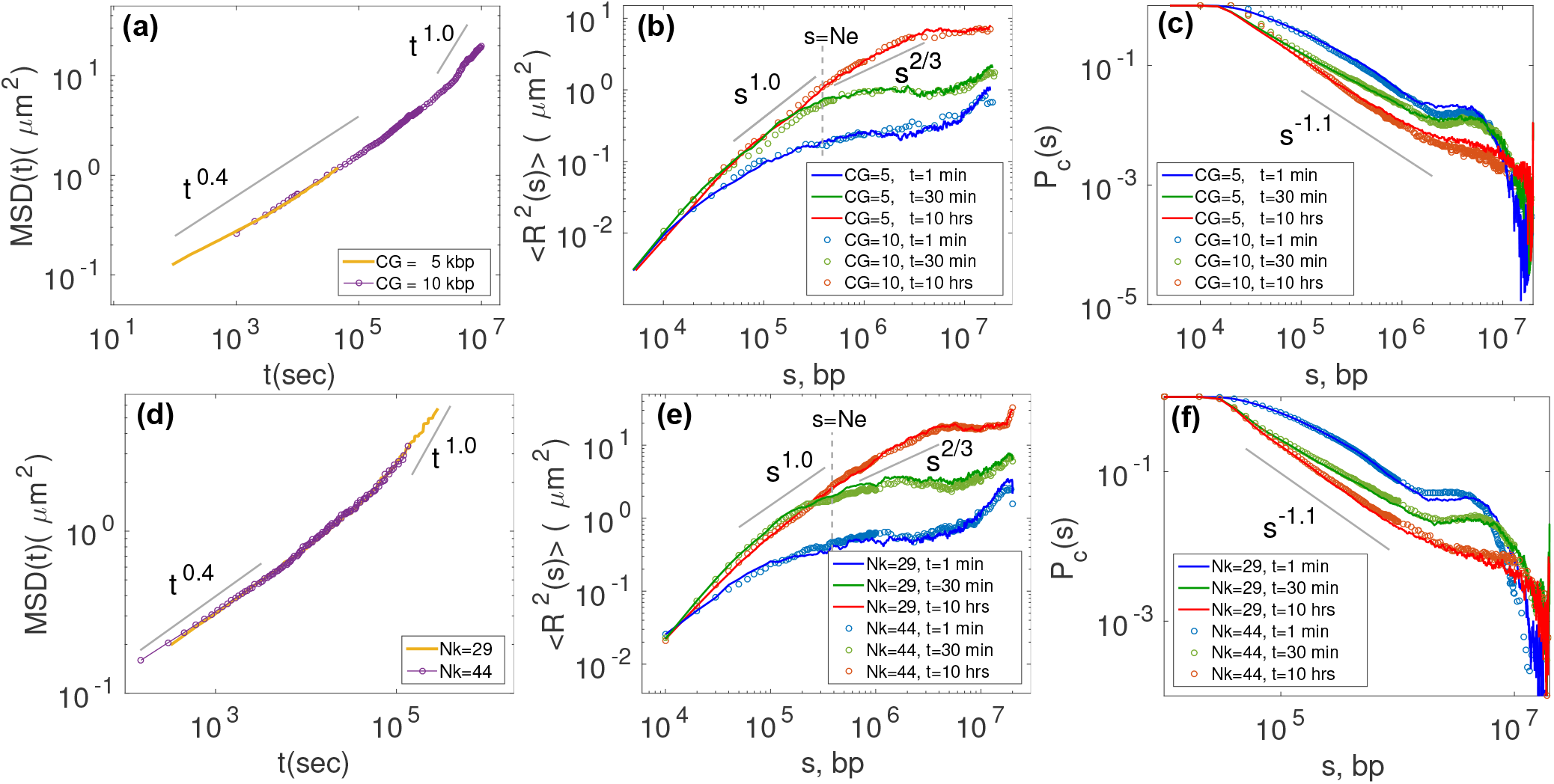
Comparison of physical properties and their time evolution for two different coarse-grainings *(CG* = 5, 10 kbp) at a fixed Kuhn size of *N_k_* = 23 kbp (top panels) or for two different Kuhn sizes (*N_k_* = 29, 44 kbp) at a fixed coarse-graining of CG=10 kbp (bottom panels) for the drosophila case *(L/L_e_* = 70, *ρ* = 0.009*bp/nm*^3^). (a,d) Average MSDs as a function of time in sec, calculated from the trajectory of 10^7^ Monte Carlo steps. Time evolution of 〈*R*^2^(s)〉 (b,e) and *P_c_(s)* (c,f) for different coarse-grainings (b,c) and Kuhn sizes (e,f).

To test that controlling the volumic fraction Φ, or equivalently the Kuhn fragment size *N_K_*, in our strategy does not impact the coarse-graining, we perform simulations at the same CG (= 10 kbp), but for different values of *N_K_* (Fig.2 and Fig.S10a,b). Identically, we observe almost perfect matching for the time-evolution of 〈*R*^2^〉, *P_c_* and *σ*^2^ *(s)* for all ranges of genomic distance. To push our strategy to the limit, we also considered very high values for Φ (Fig.S11a,b,c). In our lattice polymer model, each point can be occupied by two consecutive monomers (see Sec.4.2), so in principle, a maximum volumic fraction Φ = 2 is achievable. For Φ ≲ 1, all the simulations show exactly same results as the reference model and follow the same curve. For Φ ≳ 1 the results deviate from the reference model strongly, the dynamics are dramatically slowed down due to the incapacity of the algorithm to move the monomers efficiently.

All this demonstrates that our coarse-graining strategy allows to describe the correct structural and dynamical properties of the underlying model at all scales at steady-state but more importantly also out-of-equilibrium as long as the initial configurations share the same statistical behaviors and the chosen volumic fraction is not too high. What is the gain in term of numerical efficiency? Decreasing the number of beads by augmenting the CG would automatically linearly reduce the time to compute 1 Monte-Carlo time-step in our simulations. In addition, as the CG and the controlled Φ (or equivalently *N_K_*) are increasing, the mesh size of the lattice model (or the size of a monomer) augments, and thus 1 MCS will correspond to a larger real time step (see Tab.1). Therefore, the simulation of a fixed time period will be consequently decreased. All in all, we observed a fast polynomial decay of the numerical effort as a function of the CG scale (CPU time ~ CG^−5^, Fig.3a), with, for example, a gain of almost four orders of magnitude between the reference fine scale model and the CGM at 10 kbp resolution with *N_k_* = 23 kbp (or Φ = 0.97). Similarly we gain polynomially (CPU time ~ 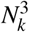) in computational time for smaller Kuhn size *N_K_* (see Fig.3b). However smaller *N_k_* corresponds to higher Φ which may impose restrictions on the dynamics if Φ > 1.

**Fig 3.**
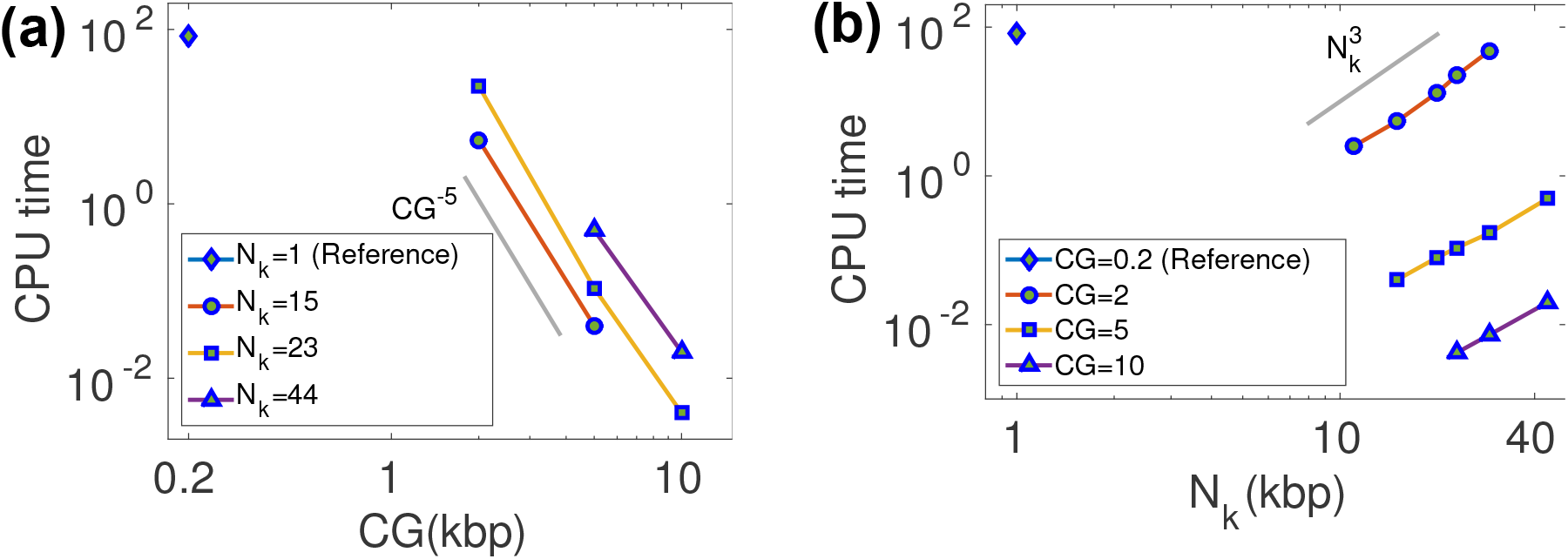
CPU time (given in hours) required to simulate 30 *min* of real dynamics for the drosophila case on a 3.20 GHz processor, (a) as a function of coarse graining where Kuhn sizes are fixed at three different values and, (b) as a function of Kuhn size at three different coarse grainings. The reference model is represented by *N_k_* = 1 kbp and *CG* = 0.2 kbp.

### 2.3 How to build a good coarse-grained null model of chromosome?

Our coarse-graining strategy is generic and has no direct connection to any particular polymer. When applied to a specific system, the question would be how to choose the optimal coarse-graining? As we observed above, regarding numerical efficiency, one wants to go for the higher CG and higher Φ (≲ 1). These two values will determine the spatial and temporal resolutions of the model. Therefore, a natural choice would be to maximize CG and Φ under the constraints of a minimal desired resolution (e.g., determined by the experimental precision). For example, for chromosomes, we aim to be quantitative typically at a 20 kbp scale (*N_K_* ~ 20 kbp) with a spatial precision of about 100 *nm.* Under this loose constraints, *CG =* 10 kbp and Φ = 0.97 is a very appropriate choice (see Tab.1).

Till now, throughout our study, we strategically choose the bending stiffness *l_k_* or the lattice volumic fraction Φ so that we preserve the physical properties of the system by conserving the right entanglement regime. Now the question is what happens if one uses more naive coarse-graining strategies that do not necessarily preserve the topological regime. As explained at the beginning of Sec.4.3, a typical strategy is to “neglect” the artificial bending rigidity if the CG is higher than the Kuhn length of the reference model. Another is to consider an isolated polymer and to neglect the “confinement” of the chain. These two kinds of models may modify the *L/L_e_* ratio and therefore may change the physical properties of the system: chain motion may be slightly accelerated (*g*_1_(*t*) ~ *t*^0.5^, Fig.4a) and structural properties may be strongly perturbed (Fig.4b,c) (see also Fig.S11 bottom and Fig.S12). In particular, considering an isolated chain (Φ ≪ 1) dramatically modifies the behavior of *P_c_* that scales within this approximation as ~ *s*^−2^, characteristics of isolated self-avoiding walks^46^. This leads to an underestimation of the contact probability by orders of magnitude compared to the reference model.

**Figure 4.**
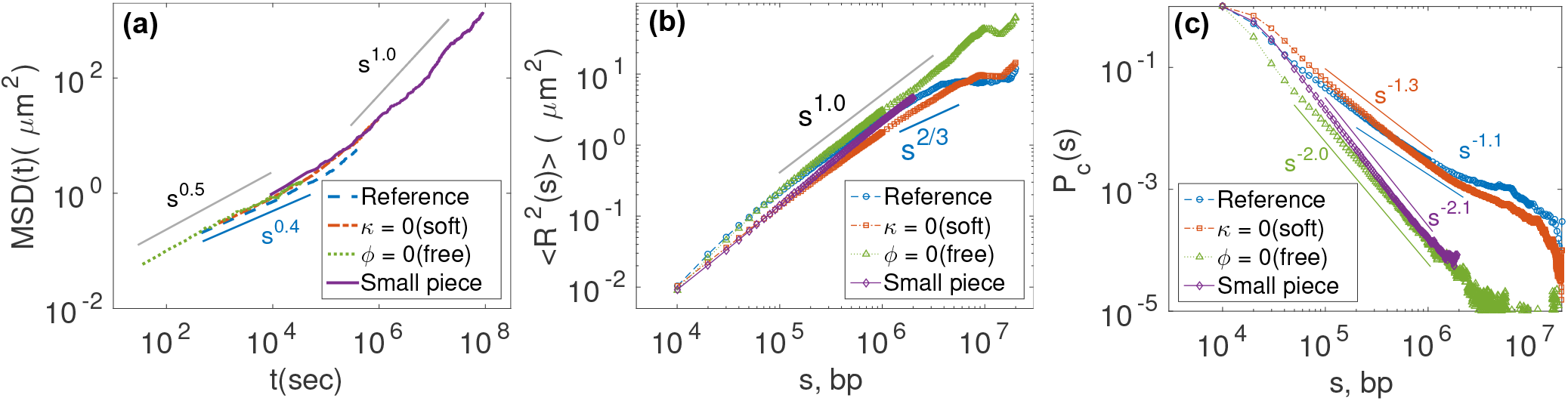
Comparison between the reference model of drosophila and various chains constructed in extreme simulation conditions: soft *k* = 0, isolated Φ << 1, and small pieces with chain length less than *N*. (a) *g*_1_ *(t)*, (b) 〈*R*^2^(*s*)〈 and (c) *P_c_*(*s*).

In complement to coarse-graining strategies, still in the purpose of reducing the computation time, a standard approach is to simulate only small pieces of the chromosomes instead of the full length. Since the dynamical regime of the chain depends on the ratio *L/L_e_*, reducing the value of *L* may modify the dynamics of the chain and therefore may lead to wrong predictions. For example, instead of 20 Mbp-long polymer, if we simulate a small fragment of 2 Mbp, we found strong discrepancies. At small length scales, *s* < 100 kbp, it follows the reference model, but at larger length scale 100 kbp < *s* < 2 Mbp it deviates from reference model and behaves like an isolated Rouse polymer (see Fig.4 and Fig.S13).

All this emphasizes the need to conserve properly the ratio *L/L_e_* of the reference model if one aims to simulate the right polymeric behavior accurately. Modifying this ratio by decreasing *L* or by making approximations that affect *L_e_* would possibly lead to simulate a system with different physical properties than the actual system of interest.

### 2.4 Application to chromatin folding in drosophila

Having in hands a strategy to build an efficient coarse-grained “null” model for chromosome, we use it to study the folding of fly chromosome 3R. In drosophila, the 3D structural units, the so-called TADs, are strongly associated with the 1D epigenomic domains^11,61,62^: a locus of a given epigenomic state is likely to share its local 3D compartment with loci of the same epigenomic state. This observed correlation had recently motivated us to build a heteropolymer model accounting for the epigenome folding into interacting TADs^20,32^. Based on biochemical evidence that proteins associated with some epigenomic states have the capacity to oligomerize^63–65^, hence possibly generating effective specific interactions between loci of the same state, we developed a block copolymer model of fly chromatin where each block represents an 1D epigenomic domain. By varying the strength of these specific interactions, we showed that such model well accounts for the TAD formation and for inter-TAD long-range contacts. Previously, we limited our analysis to short pieces of chromatin (~ 1 Mbp-long fragment) at equilibrium. In section 2.3 of the present paper, we observed that simulating only small regions instead of the full system might lead to strong approximations. Here, we wonder if our conclusions on the 3D chromosome folding in drosophila remain valid and can be generalized when considering a larger genomic region and using a more realistic “null” model for chromatin.

We consider the 20-Mbp region of chromosome 3R localized between 7 and 27 Mbp, that we model using an efficient coarse-graining (10 kbp and Φ = 0.97 for *L/L_e_* = 70, *ρ* = 0.009*bp/nm*^3^, see Table 1). For this region, we collect the epigenomic domains obtained by Filion et al^66^ for the embryonic cell line Kc167. In this dataset, five types of epigenomic states exist: 2 euchromatic states associated with active genes that, for simplicity, we decided to merge into one “active” state; and 3 heterochromatin states: constitutive heterochromatin associated with HP1-protein and H3K9me3 histone marks, facultative heterochromatin associated with Polycomb (PcG) proteins and H3K27me3 histone marks, and the so-called “black” chromatin, the prevalent form of heterochromatin. To each 10-kbp monomer of the model, we associate the corresponding epigenomic state, and we assume that monomers of the same state may specifically interact with an energy *E_i_* (in *k_B_T* unit) if they are spatially in contact (ie nearest-neighbor on the lattice)(see Materials and Methods). For simplicity, we assume that the strength of interaction is similar for every epigenomic state.

#### 2.4.1 Effect of varying the strength of specific interactions

We first concentrate on the polymer dynamics by studying the average MSD along the simulations for various values of *E_i_* (Fig.5a). For all investigated interaction energies, the scaling properties of *g*_1_ are compatible with diffusion of a crumpled polymer as seen in Sec.2.1.2 with *g*_1_ ~ *t*^0.35-0.4^. As we increase the absolute value of interaction strength, there is a dramatic slowing-down of the polymer dynamics. Interestingly, by plotting the mean MSD at 10^6^ MCS as a function of *E_i_* (Fig.5b), we observe a transition between a “fast” (*E_i_* > −0.25) and a “slow” (*E_i_* < −0.25) regime. This suggests a glass-like^67,68^ dynamic transition that occurs for strong specific interactions, reducing significantly the monomer mobility in the simulations.

**Figure 5.**
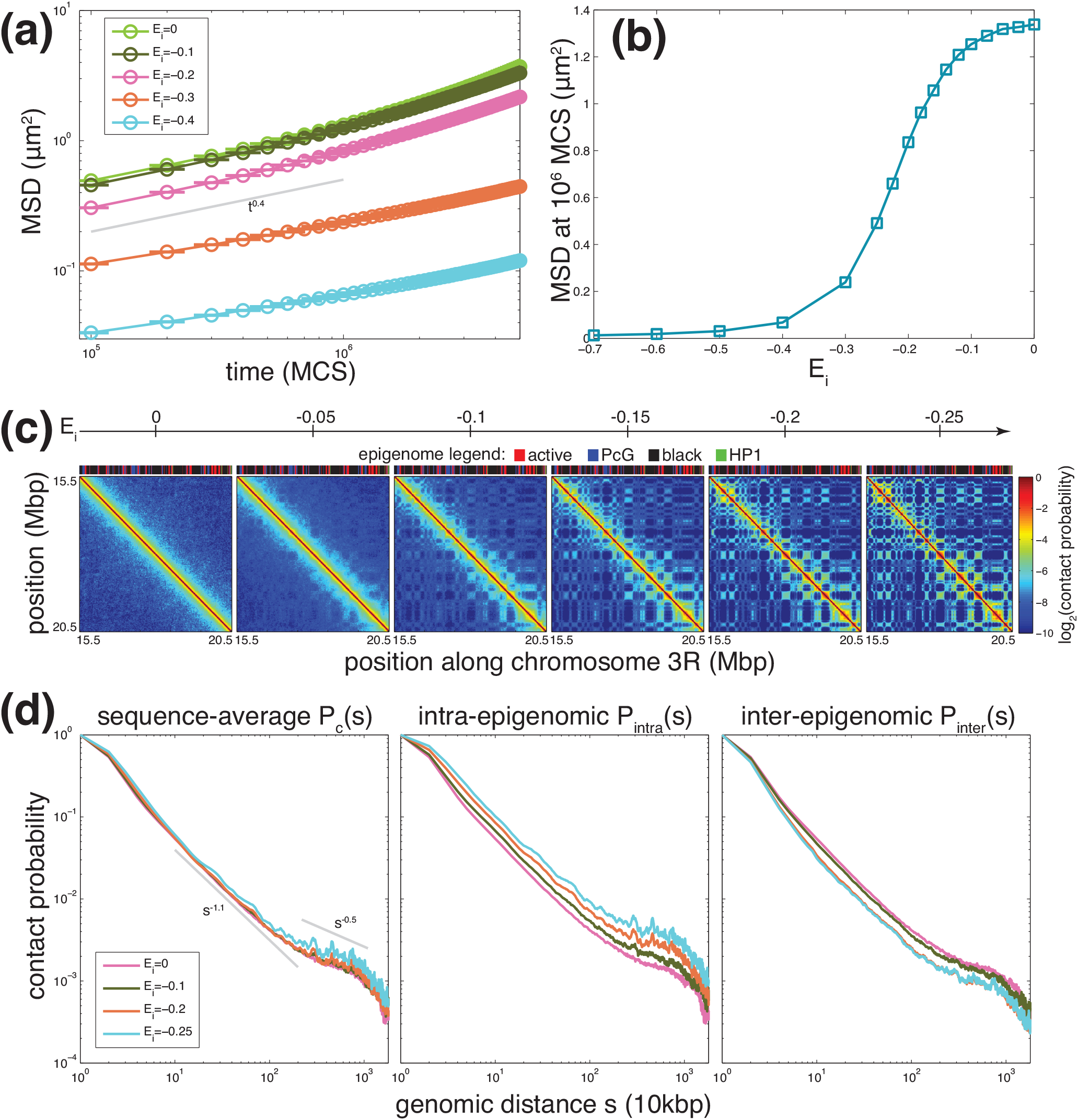
Dynamical and structural properties predicted by the model. (a) Average mean-squared displacement (MSD) along simulated trajectories as a function of simulation time in Monte Carlo step (MCS)-unit, for different values of the specific epigenomic-driven contact energy *E_i_* (in *k_B_T*-unit). (b) Average MSD after 10^6^ MCS as a function of *E_i_*. (c) Evolution of the average (between 0 and 20h) contact maps for the region located between 15.5 and 20.5 Mbp of chromosome 3R as a function of *E_i_*. We also plot on top the underlying epigenomic landscape. (d) Average contact probability as a function of the genomic distance *s* between any pairs of monomers (left), between pairs of monomers having the same (center) or different (right) epigenomic state. Grey lines represents scaling laws.

For each *E_i_*, we performed the time mapping strategy (see above) to adjust the simulation time (MCS) to the real time. Then, we computed the average contact maps between 0 and 20 hrs (*d_c_* = 163 nm), representing the average inside a population of unsynchronized cells with a typical cell cycle of 20 hrs^69^. From our 10^7^ MCS trajectories, this was possible only for *E_i_*-values in the fast regime *(E_i_* ≥ −0.25). For example, for *E_i_* = −0.3, the average map is only between 0 and 4h, and for *E_i_* = −0.4, between 0 and 300 sec. At very weak interaction strengths, the polymer is crumpled as described in Sec.2.1.2. As *E_i_* is increased, blocks (i.e. epigenomic domains), start to collapse forming TADs, long-range interactions between TAD of same type appear (Fig.5c) and TADs of different types segregate, which is characteristics of microphase separation in block copolymer^70^. From the contact maps, we estimate the sequence-average contact probability *P_c_(s)* as a function of the genomic distance s, as well as the average contact probability *Pi_ntr_α(s)* (resp. *P_in_t_er_(s))* between loci of the same (resp. different) epigenomic state. As expected, stronger interactions favor (resp. unfavor) contacts at all scales between monomers of the same (resp. different) type (Fig.5d, center and right). Interestingly, while we observe opposite behaviors for *P_intra_* and *P_inter_*, for 0 ≥ *E_i_* ≥ −0.2, the global sequence-average probability *P_c_* remains identical to the “null” model without interaction (Fig.5d, left), the increase in intra-state contacts being compensated by the decrease in inter-state contacts. At some point (*E_i_* ≤ −0.2), insulation becomes maximal and only intra-state contacts augments, leading to also an augmentation in *P_c_*.

#### 2.4.2 Comparison with experimental data

We next compare our results to Hi-C data obtained by Sexton et al for late drosophila embryos^61^. Experimental data exhibit the characteristic presence of TADs along the diagonal of the Hi-C map and of preferential long-range contacts between some TADs (Fig.6a and Fig.S14). The sequence-average probability *P_c_(s)* shows different regimes (Fig.6b): for *s* < 100 kbp, *P_c_(s)* ~ *s*^−0.5^, for 100 kbp < *s* < 1 Mbp, *P_c_(s)* ~ *s*^−1^, for 1 Mbp < *s* < 10 Mbp, *P_c_(s)* ~ *s*^−0.5^. Contacts between loci of the same epigenomic state are about 1.5-fold more pronounced at almost all scales than between loci of different states (Fig.6b).

**Figure 6.**
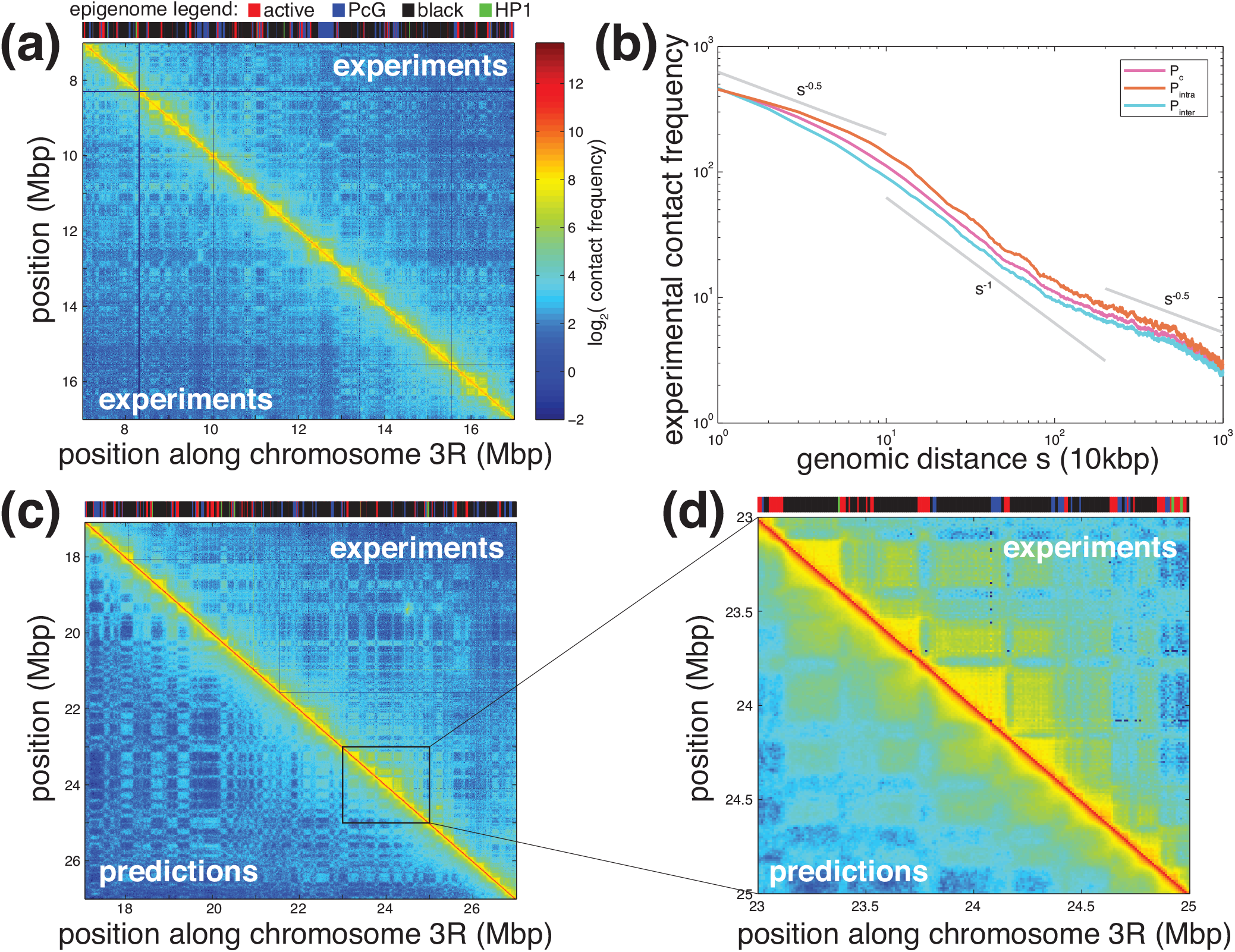
Comparison between experimental and predicted data. (a) Experimental Hi-C map for a 10 Mbp region of chromosome 3R. Corresponding epigenome is shown on top. (b) Average experimental contact frequency as a function of the genomic distance *s* between any pair of monomers (pink), between pairs of monomers having the same (orange) or different (cyan) epigenomic state. Grey lines represents scaling laws. (c, d) Predicted (*E_i_* = −0.1 *kT*) vs experimental contact maps for a 10 Mbp and a 2 Mbp region. Predicted data were multiplied by a factor 2500 to adjust scale with experiments.

While we do not expect our model to be quantitative at small genomic scales (*s* < 100 kbp) due to the coarse-graining we used in our simulations, the predicted shape of *P_c_(s)* is very similar to the experimental one with *P_c_(s)* ~ *s*^−11^ for 100 kbp < *s* < 1 Mbp and *P_c_(s)* ~ *s*^−0.5^, for 1 Mbp < *s* < 10 Mbp. Among the different strength of specific interactions that we investigated, *E_i_* = −0.1 offers the best match with experimental data (Fig.S15) with also an enrichment of 1.5 fold of intra-state vs inter-state contact probabilities. Comparison between the predicted and the experimental contact map shows very good correlations (Pearson correlation=0.86) at the local - TAD - level but also at higher scales (Fig.6c,d and Fig.S14), in terms of patterning but also in terms of relative contact frequencies. Given the simplicity of the model, it is remarkable that such model is in quantitative agreement with experimental data from small to large scales, suggesting that epigenomic-driven forces are main players of the chromosome folding in drosophila, generalizing our previous findings made on Mbp-genomic regions^20,32^.

#### 2.4.3 Dynamics of TAD formation and inter-TAD interactions

One strong conjecture of our previous study was that long-range TAD interactions are dynamical and that TADs may form very rapidly just after the mitotic exit^20^. Now that we have a more complete and largest-scale model with a proper time mapping, we aim to verify and to characterize these hypothesis. For *E_i_* = −0.1, we compute the population-average contact map of synchronized cells at different times along the simulations.

Time-evolution of the predicted Hi-C maps shows that TADs form very quickly in about half a minute (Fig.7a). Specific long-range contacts between monomers of the same epigenomic state are more slowly formed, ranging from minutes for sub-Mbp-scale contacts to hours at 10 Mbp-scale (Fig.7a). This is confirmed by analyzing the time-evolution of the average ratio between *P_intra_* and *P_c_* for different scales (Fig.7b). For 10 – 100 kbp range, *〈P_intra_/P_c_*〉 reaches a plateau after about 5 *min,* suggesting that local interactions reaches steady-state very early in the cell cycle. For the 100 kbp – 1 Mbp, convergence to steady-state is slower (less than 1 hour), while for longer-range interactions it takes more time (about 5h). Insulation between loci of different epigenomic states evolves at the same time-scales (Fig.7b).

**Figure 7.**
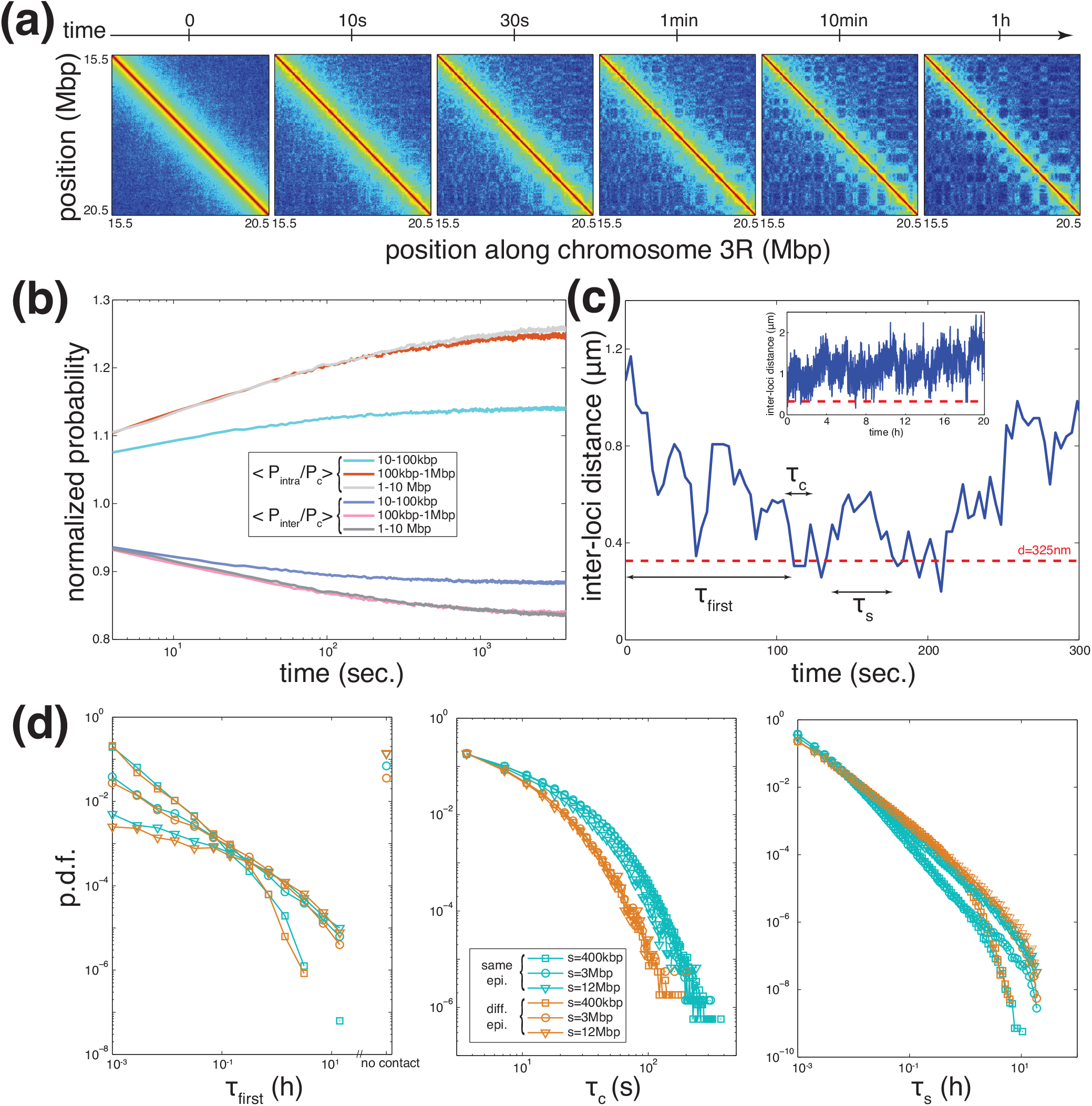
Dynamics of interactions. (a) Predicted contact maps *(E_i_* = −0.1 *kT*) for the region located between 15.5 and 20.5 Mbp of chromosome 3R as a function of time along the cell cycle. Same legend as in Fig.5c. (b) Time evolution of the ratio between *Pintrα* and *P_c_* (increasing curves) and of the ratio between *P_inter_* and *P_c_* (decreasing curves) averaged over genomic distances between 10 and 100 kbp, between 100 kbp and 1 Mbp and between 1 and 10 Mbp. (c) Example of the time evolution of distance between two loci in early times. The inset is the full trajectory along the cell cycle. The red dashed line represents the cut-off distance we choose to define that the two loci are in “contact” or not. From each trajectory, we define one value for the time of first encounter *τ_first_* and several values for the contact time *τ_c_* and the search time *τ_s_*. (d) Probability distribution functions (p.d.f) of *τ_first_* (left), *τ_c_* (center) and *τ_s_* (right) for pairs with the same (blue) or different (orange) epigenomic state separated by different genomic distance: *s* = 400 kbp (squares), 3 Mbp (circles) or 12 Mbp (triangles).

To quantify the dynamics of long-range contacts between TADs, we tracked during one cell cycle (20h) with great precision (snapshots every 100 MCS ≈ 3.5 sec). Six pairs of loci having identical or different epigenomic states (Table S3) and separated by different genomic lengths *s* (*s* ~ 400 kbp, *s* ~ 3 Mbp and *s* ~ 12 Mbp). Fig.7c shows a typical time-evolution of the distance between one pair of loci in one simulated trajectory. From the trajectories, we extract three quantities: the time of first encounter *τ_first_* defined as the first time after the mitotic exit when the pair becomes closer than *d* = 325 *nm*; the contact time *τ_c_* defined as the time the pair stays in “contact” (ie closer than *d*); and the search time *τ_s_* defined as the time interval between two “contacts”. For each pair, the probability distribution function of *τ_first_* is polynomial with two regimes with a slower decay for *τ_first_* < 0.2h (Fig.7d left). The scaling laws depend only on the genomic distance *s* between the loci, distant pairs needing more time to first contact. The polynomial dependence implies that very long *τ_first_* are significantly observed. Interestingly, for *s* > 1 Mbp, there exists a significant proportion of cells (5% for *s* ~ 3 Mbp and 14% for *s* ~ 12 Mbp) where the distance between the two regions never goes below *d*. The distributions of *τ_c_* are also polynomial for long times (Fig.7d center), the behavior depending only if pair members have the same or different epigenomic states. While, all the scaling laws are very similar, long contacts for pairs of loci with the same state are more frequent. In average, a contact between same-state loci lasts 12 seconds while the contact duration is divided by two for regions with different states. The distributions for *τ_s_* are polynomial with two regimes (Fig.7d right). While the “small” time regime depends if the epigenomic states are identical or not, different-state loci being more likely to wait more between two contacts, the “long” time regime depends mainly on the genomic scale, distant loci needing more times to contact. Indeed, for short *τ_s_*, there is still a memory of the relative positions of the two loci and pairs of the same state would be more likely to contact again rapidly, for long *τ_s_*, memory is lost and the time interval between two contacts relies only on the genomic distance as for *τ_first_*.

## 3 Discussion and Conclusion

In this article, in the first part, we introduced a new coarse-graining strategy for long and dense self-avoiding walks that conserves the entanglement length and the volumic density. Using kinetic Monte Carlo simulations on a lattice, we demonstrated that such strategy leads to the accurate description of, not only the steady-state but also the time-evolution of the expected structural and dynamical properties of the underlying fine-scale model. We showed that by introducing an effective rigidity to the coarse-grained model and by controlling the volume fraction, we could achieve very high gain in numerical (CPU-time) efficiency while maintaining a quantitative approximation and minimizing the loss in spatial and temporal resolution of the model. Using our efficient polymer model one can simulate chromosome dynamics during the whole cell cycle on a desktop computer within a day. While we illustrated our approach using chromosomes on a lattice model as toy examples, our strategy is generic and can be applied to any self-avoiding polymers and to off-lattice systems. We emphasized that neglecting topological constraints may lead to an erroneous description of the fine-scale model. Therefore the effect of supplementary interactions added to the null model, to describe specific observations present in experimental data, may lead to misinterpretation.

In the second part, we used our strategy to build a coarse-grained null model for chromatin and decorated it with a copolymer framework based solely on epigenomic data to investigate the folding and dynamics of a big fraction of chromosome 3R of drosophila. It is the first study trying to quantitatively describe the behavior and time evolution of such large genomic regions (20 Mbp) during one cell cycle (20h of real time) with high precision (10 kbp resolution). Our heteropolymer model has one unique parameter, the strength of short-range interaction *E_i_* between genomic loci having the same epigenomic state. Our findings are in qualitative agreement with our previous works based on shorter pieces of chromosomes^20,32^, but significantly improve our description of chromosome folding in drosophila. By varying *E_i_*, we showed that the system continuously switches from a dynamic homogeneous crumpled-like behavior to a crumpled heterogeneous micro-phased state. Interestingly, we observed that during this transition, the chromosome fluctuations characterized by the mean squared displacement conserve the same scaling behavior (*g*_1_ (*t*) ≈ *γt*^0.4^) with exponents compatible with a crumpled polymer. However, the prefactor *γ* depends on *E_i_* and is sharply reduced above a given strength of interaction, characteristics of a glass-like transition^67,68^. Another interesting observation was that the sequence-average contact probability *P_c_(s)*, a quantity directly comparable to experimental data, is independent of *E_i_* (at least for weak, biologically relevant values) and is same as in the reference null model, as already observed by Gursoy et al^71^. This motivates, afterward, the validity of homopolymer models, extensively used by polymer physicists, to study the generic physical principles behind chromosome folding based on comparison with sequence-average experimental data^16,41,43,58^, even if such chromatin organization is strongly heterogeneous. This also suggests that, before adding specific interactions to the system, any quantitative approach should first aim to describe such sequence-average behaviors in a null homogeneous model.

Comparing our model predictions for chromosome 3R to the corresponding Hi-C data, we observed an excellent agreement at all scales, strongly suggesting that epigenomics is a primary driver of chromosome folding in drosophila. The strength of interaction compatible with the data is weak (~ 0.1 *kT*) and locate the *in vivo* situation in the transition zone between the homogeneous crumpled and the micro-phased states. TADs are only partially collapsed and interact dynamically with other TADs of the same main epigenomic state. This suggests a substantial stochasticity in chromosome organization, consistent with recent single-cell Hi-C or super-resolution experiments^72–77^. We also detected several discrepancies between the predicted and experimental contact maps. For example, the model predicts spurious contacts or misses some between some TADs. This could be due to a wrong annotation of the local epigenomic state or the existence of specific interactions driven by other biological processes not accounted in the model. For example, refining the model to account more precisely for the local epigenetic content (for example the relative levels of histone modifications or chromatin-binding proteins) or differences in interaction strengths between different states would certainly lead to a better correspondence. We also observed that TADs are more sharply defined in the experiments, particularly in the corners of large TADs. This might be the results of the presence of cis-interacting mechanisms, like supercoiling^23,78^ or the recently proposed loop extrusion model in mammals^26,27^, that enhance the contact frequency along the genome.

To exploit the capacity of the model to simulate long trajectories (20 hrs), we analyzed the time evolution of chromosome organization. TADs formed very quickly (within minutes), entirely consistent with Hi-C data made on synchronized mammalian cells showing that, in early G1, TADs are already observable in the data^76,79^. Formation of long-range contacts require more time and eventually appear hours after the mitotic exit, also consistent with the time evolution of Hi-C data during cell cycle in human^76,79^ and yeast^80^. To go deeper into this characterization and get insights into the dynamics of contact formation, we tracked pairs of loci. At the investigated resolution (10 kbp), contacts are transient and their typical lifetime is ~ 10 seconds, indicating a very dynamic situation, consistent with many experiments performed on living cells^81^. Probability distribution functions of the first encounter time, the contact time, or the search time are polynomial, suggesting a possible connection with fractional brownian motion physics, as for bacterial chromosomes^82^. Interestingly, we predicted that a significant proportion (5 – 15%) of long-range contacts (> 1 Mbp) are not established within one cell cycle. This suggests that the genomic distance between regulatory elements should not exceed 1 Mbp to ensure that physical contact between the elements, prerequisite to an activation or repression event, would happen at least once in the cell cycle in order to maintain a stable regulation or gene expression. With the recent progress in genome editing^83^, it would be interesting to experimentally test such predictions by simultaneously tracking the distance between a promoter and its enhancer and the current gene activity^84^, for various genomic distances between the two elements. All this suggests that the 3D chromosome organization in higher eucaryotes is out-of-equilibrium and the chromatin is very dynamic. This emphasizes the necessity to properly account for the time evolution of such organization in quantitative models of chromosomes, especially for higher eucaryotes where chromosomes are strongly topologically constrained.

As a proof of concept, we demonstrated the utility of our coarse-graining approach to study chromatin organization in drosophila. However, the numerical efficiency of the method opens new perspectives to investigate the physical and mechanistic principles behind chromosome folding more deeply with many aspects remain to be understood. For example, extrapolating to the whole human genome, it would require ~ 120h of CPU time with our strategy at 10kbp resolution to simulate one cell cycle while it remains illusive to do it with the fine-scale model (> 100 years of CPU time). The possibility to easily simulate the dynamics of chromosomes or full genomes during long biologically relevant time period would allow to quantitatively investigate in the future the role of other types of interactions, like those associated with the nuclear membrane, another important player in organizing chromosomes inside the nucleus^25,85^ and the crosstalk with epigenomic-driven interaction as presented here. This situation seems particularly attractive to describe the reorganization of chromatin during senescence where constitutive heterochromatin detaches from the membrane to form large foci at the interior of the nucleus^86^.

## 4 Materials and Methods

### 4.1 Estimation of the chromatin volumic density

The chromatin volumic density *ρ* is defined as the ratio between the genome size *G* and the average volume *V* of the nucleus. For haploid yeast, *G* = 12.2*Mbp* and *V* ≈ 2.6*μm*^3^^69,87^, leading to *ρ* ≈ 0.005 *bp/nm*^3^. For drosophila (diploid) late embryos, nuclei have a typical diameter of 4*μm*^88^ and contains about 300*Mbp* of genomic DNA, thus *ρ* ≈ 0.009 *bp/nm*^3^. In mammals, the size of the nucleus depends strongly on the cell type typically ranging from 5 to 15μm in diameter^69^. For example, for a human nucleus (*G* ≈ 6*Gbp*) of diameter 9*μm*, *ρ* ≈ 0.015 *bp/nm*^3^. For larger nuclei (12*μm* in diameter), as measured by Muller et al^89^, *ρ* may be weaker (≈ 0.007 *bp/nm*^3^).

### 4.2 Simulation of structural and dynamical properties of the null model

The polymer is modeled as a semi-flexible self-avoiding walk, consisting of *N* beads, on a face centered cubic (fcc) lattice of size *S × S × S* (each unit cell contains 4 lattice sites) following the model developed by Hugouvieux et al^90^ (Fig.8) (more details can be found in^32,90,91^). As in the elastic lattice model introduced recently by Schram and Barkema^92^, we authorize at maximum two monomers to occupy the same lattice site if and only if they are consecutive along the chain^90,92^ (Fig.8). Otherwise, due to excluded volume, two non-consecutive monomers cannot be located at the same site. When two successive monomers occupy a lattice site, an extra bond length is accumulated in that node as a ‘stored length’^92–94^ (Fig. S16b). The concept of stored length was first introduced by Rubinstein in his pioneering article on the implementation of repton model on a lattice^93^. Such double occupancy of consecutive monomers accounts for the effect of contour length fluctuations^94^. Rigidity is accounted using a standard formulation^95^:

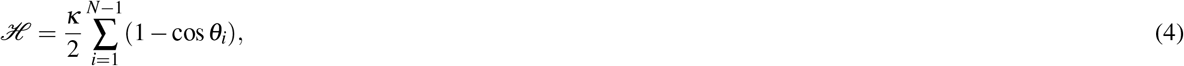

where *κ* is the bending rigidity and is directly related to *l_k_/b* (see below), and *θ_i_* is the angle between the bond vectors *i* and *i* + 1. Confinement and effect of other chains are approximated using periodic boundary conditions, the corresponding lattice volume fraction being Φ = *N*/ (4*S*^3^). Note that such periodic conditions do not confine the chain to the finite volume of the simulation box. Using correct unfolded coordinates, chains can extend over any large distances.

**Figure 8.**
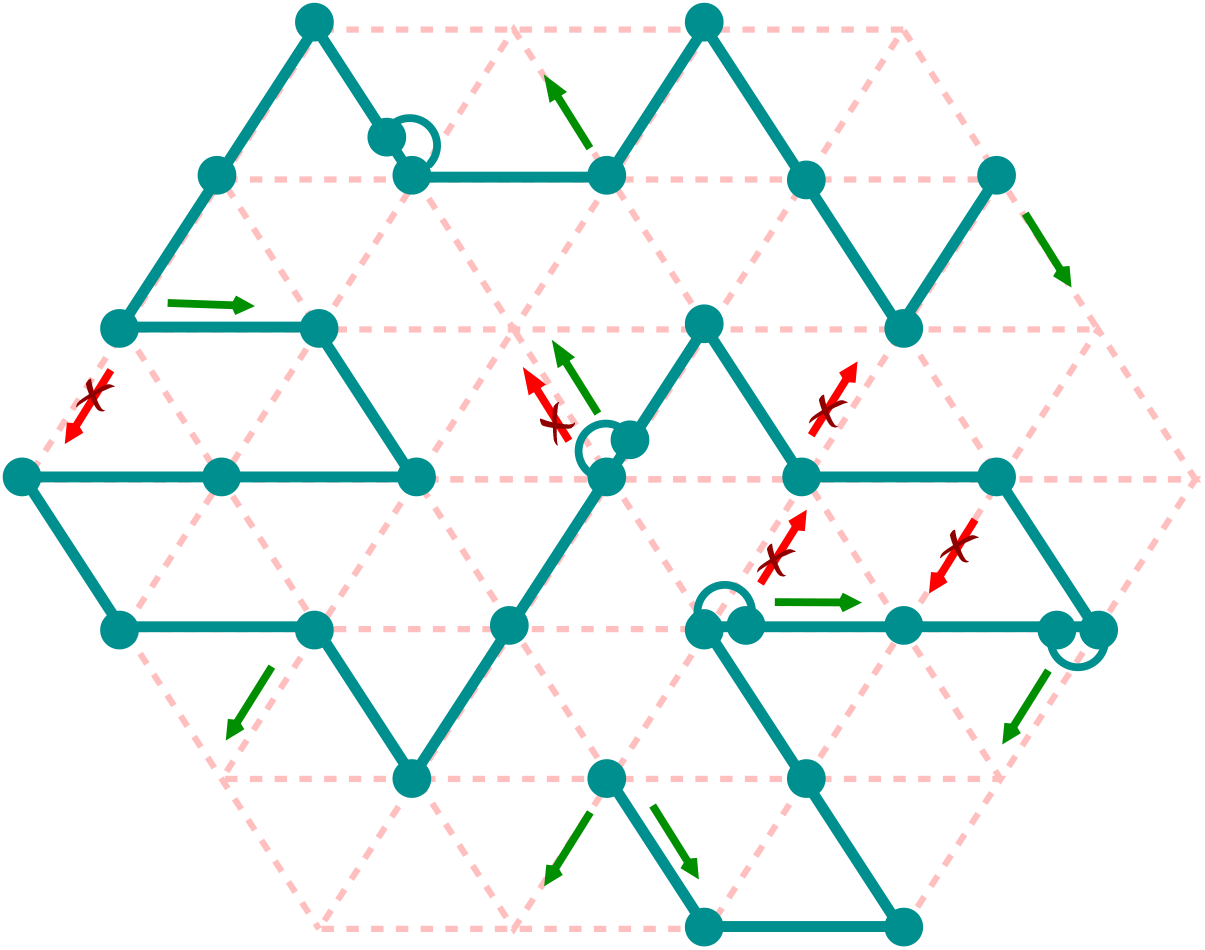
Schematic of a lattice polymer configuration on a 2D projection of a 3D fcc lattice. Solid line with beads represents the polymer chain, and the dotted lines represent the underlying lattice. Each lattice site is allowed to contain a maximum of two beads if and only if they are consecutive to each other along the chain. Semicircular arcs indicate doubly occupied lattice sites. Some of the allowed and forbidden moves are shown in green and red respectively.

The dynamics of the chain follows a kinetic Monte-Carlo (KMC) scheme with simple local moves^90^: one Monte Carlo step (MCS) consists of *N* trial moves where a monomer is randomly chosen and randomly displaced to a nearest-neighbor site on the lattice (Fig.8). Trial moves are accepted according to a Metropolis criterion applied to 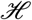 and if the chain connectivity is maintained. Compared to standard Monte Carlo methods used to study systems at thermal equilibrium^96^, KMC has the advantage to track the equilibrium or out-of-equilibrium dynamical evolution of a system. The transition rates in the KMC are assumed to be Poissonian which is likely to be an approximation of the exact dynamics at the relevant time scales. However, the connection between the real-time and KMC steps can be established precisely within the framework of Poissonian processes^97^. In our model, due to the approximated transition rates, accuracy is not guaranteed for time-scales below few MCS (temporal resolution) and for length-scales below few *b* (spatial resolution).

This KMC scheme coupled to the notion of stored length (see above) allows efficient simulations of reptation motion in dense - topologically constrained - systems, while still accounting for the main characteristics of polymer dynamics like polymer connectivity, excluded volume, and non-crossability of polymer strands^92,93^. Due to the simplicity and efficiency of such frameworks, they have been widely used in the literature to investigate various properties of polymeric systems^32,90–92,94,98–100^.

In the entangled regime (*L/L_e_* ≫ 1), dynamics could be very slow and the system may keep the “memory” of its initial configuration and topology for a long time. Chromosomes are thought to be mostly knotted-free structures^16,43,101^. Therefore, we initiate our simulations by knotted-free configurations generated using the “hedgehog” algorithm^59,102,103^: starting from a central unknotted scaffold, configurations are iteratively grown by randomly inserting monomers at nearest-neighbor sites common to two already placed consecutive monomers (Fig.S1a). We verified that, starting from other type of unknotted configurations such as Rosette(Fig.S1e) and Cylindrical(Fig.S1f). We recovered the same scaling laws for the null model (Fig.S3).

Starting from a given initial configuration, we then normally simulate 10^7^ - 10^8^ MCS and store the configurations after each 10^3^ MCS. In some special cases where we are specifically interested in small time scales, we collect configurations more frequently. From these snapshots taken from 10^2^ simulated trajectories, we then estimate several structural and dynamical quantities of interest. We focus on the time-evolution of the mean squared displacements (MSD) of individual monomers (*g*_1_ (*t*)) and of the center of mass of the chain (*g*_3_(*t*)), as well as the average squared distance 〈*R*^2^(*s*)〉 and contact probability *P_c_*(*s*) between monomers separated by a linear distance *s* along the chain. For the latter, a contact is defined if the physical distance between a pair of genomic loci is less than a particular distance *d_c_*. Note that all such properties are estimated using the ‘unfolded’ polymer conformations.

### 4.3 Relation between Kuhn length *l_K_* and lattice parameters

In this section we derived the relation between Kuhn length *l_K_* and different lattice parameters, expressed in Eq.3. We consider an arbitrary fine-scale model (FSM) of a semi-flexible self-avoiding walk defined by 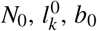. We note *N*, *l_k_* and *b*, the corresponding values of a coarse-grained model (CGM) of the FSM. Each CGM monomer encompasses *n* = *N*_0_/*N* > 1 FSM monomers.

Using Eq.1, conservation of *L/L_e_* and conservation of the volumic density in FSM monomer *ρ_FS_* leads to

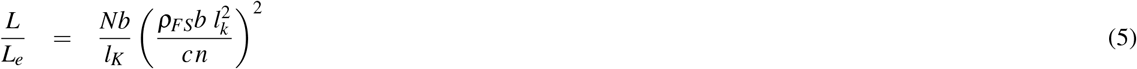

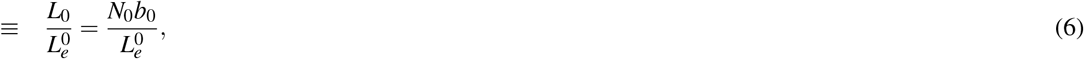

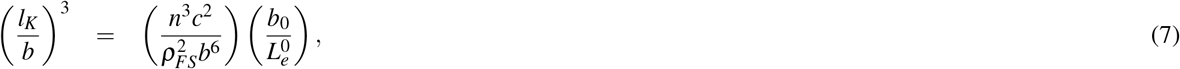

*ρ_FS_* = (*N*_0_/*V*)(= *ρ/ν*_0_ in the context of chromosome) with *V* the volume of the box. Conservation of the volume implies that

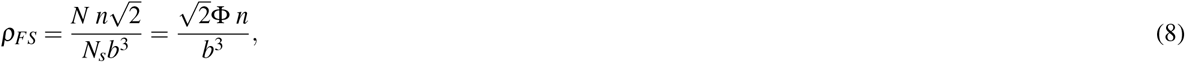

with *N_s_* the number of lattice sites and Φ = *N/N_s_* the lattice volumic fraction. Incorporating Eq.8 into Eq.7 leads to

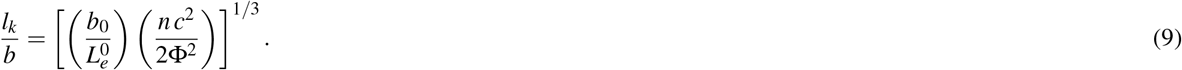

Practically, knowing *ρ_FS_*, *b*_0_ and 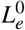 (from Eq. 1 of the main text) for the reference model, for a given coarse-graining (defined by n), Eq.9 gives us a relation between *l_k_/b* (which is related to the bending energy of the model, see below) and the lattice volumic fraction Φ. Eq.8 is used to compute the corresponding value for *b*.

### 4.4 Relation between Kuhn size *N_k_* and bending rigidity *κ* for lattice polymer

For a lattice phantom chain with *N* beads, the mean squared end-to-end distance 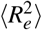 is given by^104^

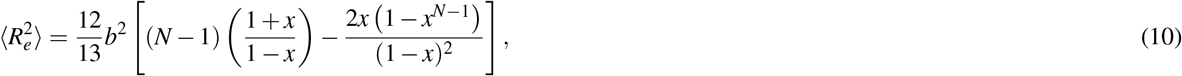

where

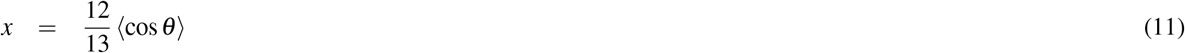

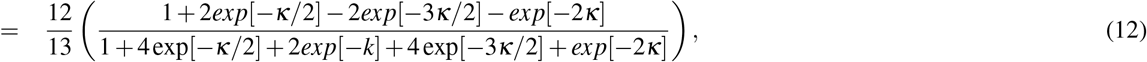

where *θ* is the angle between two monomers and *κ* is the bending rigidity correspond to the bending energy *E* (*θ*) = *κ*(1 - cos *θ*). Now we have *l_K_/b* = (1 + *x*)/(1 – *x*), which relates *N_K_* to *κ*.

### 4.5 Simulation of the block copolymer model for drosophila

In the block copolymer model, the energy of a given configuration is given by

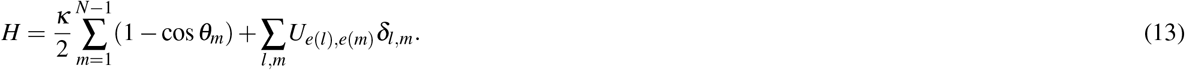

The first contribution accounts for the null model. The second contribution accounts for epigenomic-driven interactions with *δ_lm_* = 1 if monomers *l* and *m* occupy nearest-neighbor (NN) sites on the lattice (*δ_lm_* = 0 otherwise), *e*(*l*) the epigenomic state of monomer *l* and *U_e,e′_* the strength of interaction between a pair of spatially neighbor beads of epigenomic states *e* and *e*′. For simplicity, we will assume that interactions occur only between monomers of the same chromatin state (*U_e, e′_* = 0 if *e* ≠ *e*′) and that the strength of interaction (that we note *E_i_*) is the same whatever the chromatin state (*U_e,e_* ≡ *E_i_* for all *e*). Dynamics of the chain follows the same KMC scheme as the null model using a Metropolis criterion applied to *H*. For various values of *E_i_*, we simulate 400 trajectories during 10^7^ MCS starting from a random unknotted “hedgehog” configuration (as in Fig.S1a,c) at a standard *in vivo* bp-density (*ρ* = 0.009*bp/nm*^3^) (see the Movie for example). Note that such initial configurations might not reflect exactly the post-mitotic organization of fly chromosome and may impact the very large-scale - out-of-equilibrium - behavior predicted by the model.

## Supporting information

**S1 Text:** A single pdf file containing 16 supporting figures and 3 supplementary tables.

**S1 Video:** Time evolution of random unknotted ‘hedgehog’ configuration with specific interactions.

## Acknowledgements

We thank Magali Richard, Ivan Junier and Cedric Vaillant for critical reading of the manuscript, as well as Ralf Everaers, Pascal Carrivain, Aurelien Bancaud, Mikhail Tamm and Giacomo Cavalli for fruitful discussions. We acknowledge the CIMENT infrastructure (supported by the Rhone-Alpes region, Grant CPER07_13 CIRA) for computing resources. This work was supported by Agence Nationale de la Recherche (ANR-15-CE12-0006 EpiDevoMath), Fondation pour la Recherche Medicale (DEI20151234396) and CNRS.

